# Translational validation of shotgun proteomics findings in cerebrospinal fluid of sporadic cerebral amyloid angiopathy patients

**DOI:** 10.1101/2024.01.15.575618

**Authors:** Marc Vervuurt, Anna M. de Kort, Iris Kersten, Astrid van Rens, Catharina J. M. Klijn, Floris H. B. M. Schreuder, Dirk J. Lefeber, H. Bea Kuiperij, Marcel M. Verbeek

## Abstract

**Background:** Prior research conducted in model rats of CAA Type 1 (rTg-DI) identified a range of cerebrospinal fluid biomarker candidates associated with sCAA pathology. This list of potential biomarkers includes the lysosomal proteases cathepsins B and S (CTSB/CTSS) and hexosaminidase B (HEXB). It is yet unknown if these findings obtained in rTg-DI rats translate to differential protein levels and/or enzyme activities in cerebrospinal fluid (CSF) of sCAA patients. In this study, we attempted to validate CTSB, CTSS and HEXB in CSF as potential biomarkers for sCAA in a human population.

**Materials and methods:** We have included sCAA patients (n = 34) and control participants (n = 27) from our BIONIC/CAFE cohort. We analysed the CSF of these participants with ELISA for protein levels of CTSB and CTSS. Additionally, we used in-house enzyme assays to determine activity levels of total hexosaminidase and hexosaminidase A (HEXA) in CSF. The proportion of HEXA activity to total HEX activity was used as a proxy for HEXB activity.

**Results:** CSF CTSB and CTSS protein levels were not significantly different between sCAA and controls (p = 0.21 and p = 0.34). Total HEX activity was unaltered as well (p = 0.11), whereas a significant decrease was observed in HEXA activity levels (p = 0.05). HEXA / total HEX activity levels (as a proxy for HEXB activity) were unaltered between sCAA patients and controls (p = 0.19). Additionally, CTSB and CTSS protein levels positively associated with total HEX activity (r_sp_ = 0.37, p = 0.005; r_sp_ = 0.40, p = 0.003).

**Conclusion:** The contrasting results between biomarker discovery in rats and validation in human participants highlight the challenges and complexities of biomarker research. These findings offer valuable insights into the nuances of disease and the difficulties in translating laboratory findings using animal models to clinical practice. Understanding these discrepancies is essential for improving the precision of biomarker translation, ensuring clinical relevance, and developing comprehensive biomarker panels for CAA and related conditions.

## Introduction

Cerebral amyloid angiopathy (CAA) is a complex neurovascular disorder that presents a daunting diagnostical challenge to clinicians (1). Characterized by the accumulation of amyloid-beta peptides within the walls of cerebral blood vessels, CAA has been strongly linked to a spectrum of cognitive and neurological impairments, including intracerebral haemorrhage, transient focal neurological episode and cognitive impairment (2). To decipher the intricate molecular mechanisms underlying CAA, previous research focused on exploring the proteomic landscape of cerebrospinal fluid (CSF) in rat model of microvascular CAA, the rTg-DI model (3–5). Through use of data-independent acquisition liquid-chromatography tandem-mass spectrometry (DIA LC-MS/MS) proteomics, we have identified a range of potential biomarkers in rats which might be capable of differentiating patients with CAA from control subjects. Among these candidates, increased levels of CSF cathepsin B (CTSB), cathepsin S (CTSS), and hexosaminidase B (HEXB) were observed. These proteases were also previously identified as upregulated in cerebrovascular tissue of rTg-DI rats (6–8).

CTSB and CTSS, both lysosomal cysteine proteases, are integral to processes such as protein degradation, antigen presentation, and immune regulation(9). In recent years, microglial cathepsins have been implied to be involved in neuroinflammatory processes, observed in various forms of neurodegeneration, like Alzheimer’s disease (AD) (10–13). In line with this reasoning, CTSB has been considered as potential therapeutic target for neurodegenerative conditions like Alzheimer’s disease, presenting the opportunity to modulate its activity for therapeutic benefit in these contexts (14). Hexosaminidase A (HEXA), B (HEXB) and S (HEXS) are members of the hexosaminidase enzyme family, characterized by their roles in glycosphingolipid metabolism (15). More specifically, HEXA and HEXB work in concert to degrade N-acetylglucosamine (GlcNAc) and N-acetylgalactosamine (GalNAc) containing glycosylation motifs, for example on glycosphingolipid GM2 gangliosides. Genetic defects in HEXA and HEXB include the gangliosidoses Sandhoff and Tay-Sachs disease, which are characterized by negligible total hexosaminidase and hexosaminidase A activities respectively (16). Although Sandhoff and Tay-Sachs disease are unrelated to CAA pathology, gangliosides have also been implied to be of effect in Aβ-related neurodegenerative processes (17).

In this research, we applied targeted protein assays to validate the biomarker status of CTSB and CTSS (using ELISA), and HEXB (using an in-house enzyme activity assay) as biomarkers for sCAA.

## Materials and methods

### Patient inclusion

We have included patients with sporadic CAA (sCAA) and age- and sex-matched control participants in this study. sCAA patients were included from the Radboud University Medical Center as part of the BIONIC/CAFE (“BIOmarkers for cogNitive Impairment due to Cerebral amyloid angiopathy/Cerebral Amyloid angiopathy Fluid biomarkers Evaluation, www.radboudumc.nl/BCS) studies. The BIONIC/CAFE studies were cross-sectional cohort studies on new body fluid biomarkers for sCAA. These studies were approved by the Medical Ethics Committee Arnhem-Nijmegen 2017-3810 (BIONIC) and 2017-3605 (CAFE). Informed consent was collected from all participants.

In short, the major inclusion criterion for patients with CAA was a diagnosis of probable CAA according to the modified Boston Criteria; last symptomatic intracerebral haemorrhage had to be at least 3 months before inclusion (18). The inclusion criteria for the control subjects included no history of cognitive impairment, dementia, stroke or major brain pathology. Exclusion criteria for both patients with sCAA and controls included contra-indications for lumbar puncture or 3 Tesla brain MRI. All sCAA participants and control subjects included from the BIONIC/CAFE studies underwent a comprehensive assessment that included clinical and neuropsychological tests (including the Montreal Cognitive Assessment or MOCA) venepuncture- and lumbar punctures (19). Structural magnetic resonance imaging (3.0 Tesla MRI scan (Siemens Magnetom Prisma, Siemens Healthineers, Erlangen, Germany) was also conducted and included T1, T2, FLAIR and SWI sequences. The following small vessel disease markers were assessed: number of lobar microbleeds, degree of cortical superficial siderosis, degree of white matter hyperintensities, and the extent of enlarged perivascular spaces in the centrum semi-ovale (20). These findings were combined to create a CAA small-vessel disease burden score (CAA SVD).

Structural magnetic resonance imaging was also conducted to assess brain characteristics such as the number of microbleeds, degree of cortical superficial siderosis (ranging from 0 to 2), degree of white matter hyperintensities (Fazekas score ranging from 0 to 1), and the extent of enlarged perivascular spaces (scored from 0 to 1). These findings were combined to create a CAA small-vessel disease burden score (CAA SVD), which ranged from 0 to 6 (21). Further information on the inclusion- and exclusion criteria of all subjects can be found in (20).

sCAA patients (n = 34) and control subjects (n = 27) were age- and sex matched (p = 0.60 and p = 0.61) [**Table 1**]. CSF biomarkers showed a typical CAA profile, with decreased levels of CSF Aβ40 (p < 0.001) and Aβ42 (p < 0.001), but normal total tau (t-tau; p = 0.12) and threonine-181 phosphorylated tau levels (p-tau; p = 0.12).

**Table 1:**
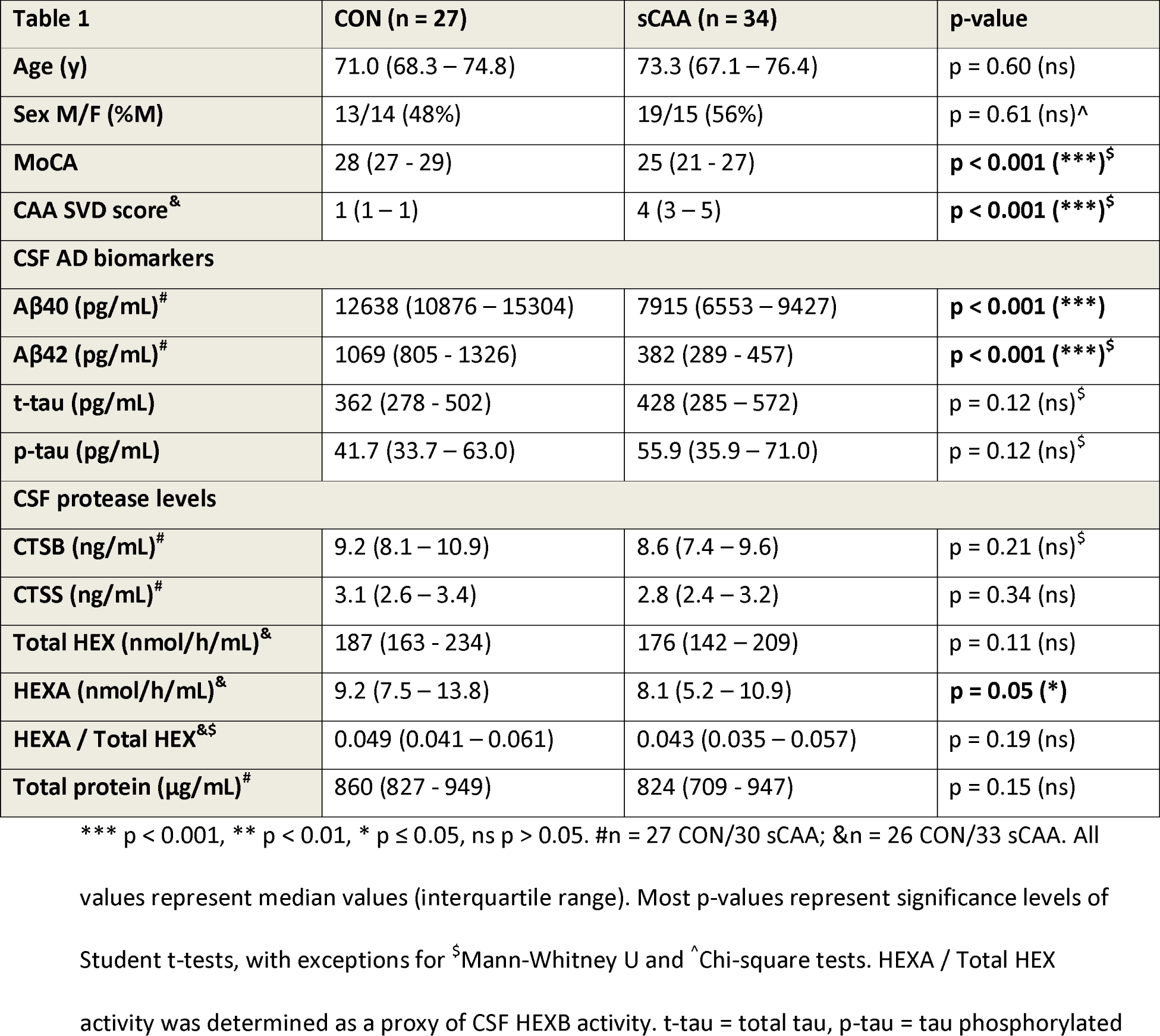
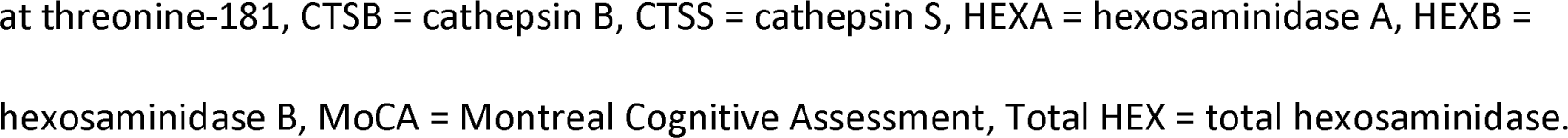
Subject characteristics per patient/control group.

| Table 1 | CON (n = 27) | sCAA (n = 34) | p-value |
| --- | --- | --- | --- |
| Age (y) | 71.0 (68.3 – 74.8) | 73.3 (67.1 – 76.4) | p = 0.60 (ns) |
| Sex M/F (%M) | 13/14 (48%) | 19/15 (56%) | p = 0.61 (ns)<sup>^</sup> |
| MoCA | 28 (27 - 29) | 25 (21 - 27) | p < 0.001 (***)<sup>§</sup> |
| CAA SVD score<sup>&</sup> | 1 (1 – 1) | 4 (3 – 5) | p < 0.001 (***)<sup>§</sup> |
| CSF AD biomarkers |  |  |  |
| Aβ40 (pg/mL)<sup>#</sup> | 12638 (10876 – 15304) | 7915 (6553 – 9427) | p < 0.001 (***) |
| Aβ42 (pg/mL)<sup>#</sup> | 1069 (805 - 1326) | 382 (289 - 457) | p < 0.001 (***)<sup>§</sup> |
| t-tau (pg/mL) | 362 (278 - 502) | 428 (285 – 572) | p = 0.12 (ns)<sup>§</sup> |
| p-tau (pg/mL) | 41.7 (33.7 – 63.0) | 55.9 (35.9 – 71.0) | p = 0.12 (ns)<sup>§</sup> |
| CSF protease levels |  |  |  |
| CTSB (ng/mL)<sup>#</sup> | 9.2 (8.1 – 10.9) | 8.6 (7.4 – 9.6) | p = 0.21 (ns)<sup>§</sup> |
| CTSS (ng/mL)<sup>#</sup> | 3.1 (2.6 – 3.4) | 2.8 (2.4 – 3.2) | p = 0.34 (ns) |
| Total HEX (nmol/h/mL)<sup>&</sup> | 187 (163 - 234) | 176 (142 – 209) | p = 0.11 (ns) |
| HEXA (nmol/h/mL)<sup>&</sup> | 9.2 (7.5 – 13.8) | 8.1 (5.2 – 10.9) | p = 0.05 (*) |
| HEXA / Total HEX<sup>&§</sup> | 0.049 (0.041 – 0.061) | 0.043 (0.035 – 0.057) | p = 0.19 (ns) |
| Total protein (μg/mL)<sup>#</sup> | 860 (827 - 949) | 824 (709 - 947) | p = 0.15 (ns) |

\*\*\* p < 0.001, \*\* p < 0.01, \* p ≤ 0.05, ns p > 0.05. #n = 27 CON/30 sCAA; &n = 26 CON/33 sCAA. All
values represent median values (interquartile range). Most p-values represent significance levels of
Student t-tests, with exceptions for <sup>§</sup>Mann-Whitney U and <sup>^</sup>Chi-square tests. HEXA / Total HEX
activity was determined as a proxy of CSF HEXB activity. t-tau = total tau, p-tau = tau phosphorylated
at threonine-181, CTSB = cathepsin B, CTSS = cathepsin S, HEXA = hexosaminidase A, HEXB = hexosaminidase B, MoCA = Montreal Cognitive Assessment, Total HEX = total hexosaminidase.

### CSF collection and biomarker assays

Every participant underwent a lumbar puncture according to standard procedures, to collect CSF. CSF was collected in polypropylene tubes, centrifuged, aliquoted, and subsequently stored at −80°C.

ELISAs were used to quantify CTSB and CTSS levels in CSF (both R&D DuoSets, R&D Systems, Minneapolis, USA). CSF samples were diluted 20x with reagent diluent prior to ELISA analysis. Additionally, we attempted to develop in-house assays for the determination of CTSB and CTSS activity in CSF (22, 23). For CTSB, 50 μL CSF samples were diluted 1:1 with 200 mM sodium phosphate, 10 mM disodium EDTA, 10 mM dithiothreitol assay buffer (pH 7.5). and incubated for 5 minutes at 37°C. Subsequently, 50 μL of 25 μM Z-RR-AMC substrate (Enzo Life Sciences, Farmingdale, USA) was added. Fluorescence was measured at 365/470 nm for a duration of 60 minutes, with interval measurements every 5 minutes. For CTSS, a similar protocol was employed: CSF samples were similarly diluted as for CTSB, after which samples were incubated for 60 minutes at 37°C. This incubation step inhibits the enzymatic activity of Cathepsin L, a cathepsin with similar substrate selectivity as CTSS. Following the incubation, 50 μL of 25 μM Z-VVR-AMC (Enzo Life Sciences, Farmingdale, USA) was diluted in assay buffer and added to the sample, after which fluorescence was measured at 380/460 nm for a total of 60 minutes, every 5 minutes.

While generic substrates are available for all hexosaminidase subunits, specific substrates only exist for HEXA. Similar substrates to which HEXB is specific are not commercially available. As the majority of total HEX activity is generated by HEXB, the ratio between specific HEXA and generic total HEX activity was used as a proxy of HEXB activity. This was done using in-house fluorometric enzyme activity assays. In these assays, CSF was diluted 7x with diluent, after which it was incubated with 12.5 mM 4-methylumbelliferyl-N-acetyl-β-D-glucosaminide (total HEX; Sigma Aldrich, MS, USA) or 5 mM 4-methylumbelliferyl N-acetyl-β- D-glucosaminide-6-sulfate (HEXA; Sigma Aldrich, MS, USA) in 0.1 M citric acid/NaOH buffer (pH 4.4) substrate buffer (24). Following 10 minutes of incubation at 37°C, the reaction was stopped using 0.5 M glycine buffer + 0.025% Triton X-100 (pH 10.6). Fluorescence was measured on a Victor2^TM^ D Fluorometer (Perkin Elmer, Waltham, USA) at (excitation/emission) = 365/460 nm.

Total protein levels of all CSF samples were determined by use of a Pierce^TM^ BCA protein assay kit (Thermo Fisher Scientific, Waltham, USA). CSF Aβ40/Aβ42 and total tau (t-tau)/tau phosphorylated at threonine-181 (p-tau) levels were determined using Lumipulse chemiluminescent immunoassays (Fujirebio, Tokyo, Japan). All standards, controls and samples were assayed in duplicate in all assays.

### Statistical analyses

We analysed the data using IBM SPSS Statistics Version 25.0.0.1 (IBM Corp., Armonk, USA) and Graphpad Prism version 5.03 (Graphpad Software, Inc., San Diego, USA).

Shapiro-Wilk tests were used to assess normality of data. Correlations between variables were determined by use of Spearman correlation analyses. Differences between groups were determined using Student’s t-tests (for parametric data) or Mann-Whitney U tests (for non-parametric data). Also, sex differences between groups were determined using Chi-squared tests. Corrections for the potential confounding (residual) influences of age and/or CSF total protein levels were made according to linear regression modelling. Test results were deemed statistically significant with a p-value ≤ 0.05.

## Results

CSF levels of CTSB and CTSS were not significantly different in sCAA and control subjects (p = 0.21 and p = 0.34 respectively) [**Figure 1 A-B**], also after correction for age (CTSB, p = 0.54; CTSS, p = 0.27). Correction for the influence of total protein levels on CTSB (p = 0.81) and CTSS (p = 0.61) did not improve significance of results. This was also the case for correction for both age and total protein levels for CTSB (p = 0.80) and CTSS (p = 0.52). Unfortunately, we were unable to optimize the CSF CTSB and CTSS activity assays in such a way that we were able to reach the necessary analytical sensitivity for a reliable and robust assay.

**Figure 1:**
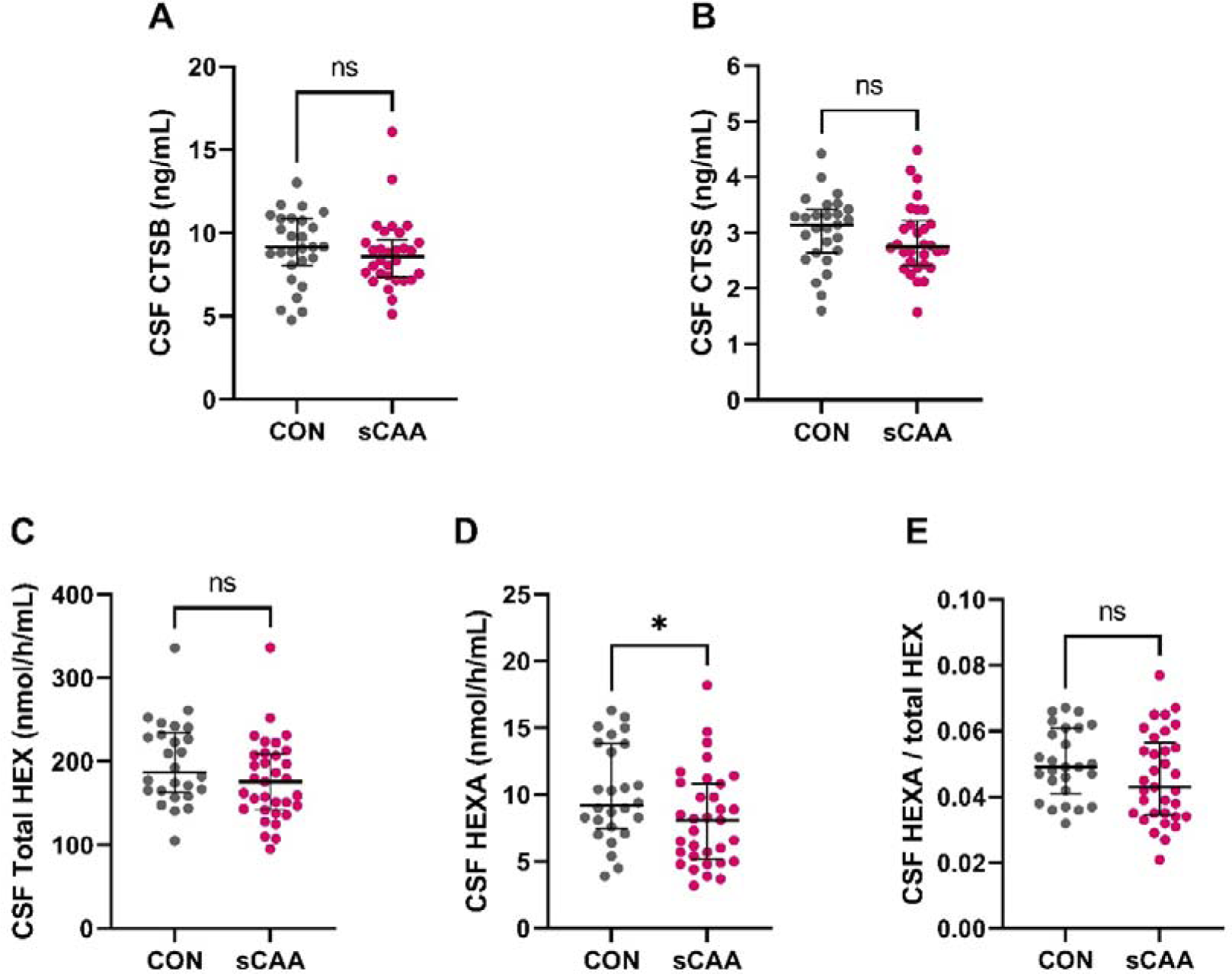
Dot plots on CSF protein levels of CTSB/CTSS and CSF activity levels of total HEX and HEXA. **(A)** CSF CTSB levels in sCAA patients and control subjects. **(B)** CSF CTSS levels in sCAA patients and control subjects. **(C)** CSF total hexosaminidase activity (HEX) in sCAA patients and control subjects. **(D)** CSF hexosaminidase A (HEXA) activity in sCAA patients and control subjects. **(E)** CSF HEXA / total HEX activity, as a proxy for CSF HEXB activity, in sCAA patients and control subjects. *p < 0.05, ns = not significant

The enzyme activity assay for total hexosaminidase activity also revealed no significant differences between sCAA patients and control subjects (p = 0.11). HEXA activity levels were significantly decreased in sCAA participants compared to controls (p = 0.05) [**Figure 1 C-D**]. Correcting total HEX and HEXA activity levels for age (p = 0.11; p = 0.05, not significant), total protein (p = 0.20; p = 0.10) or both age and total protein (p = 0.19; p = 0.11) did not produce significant differences between sCAA and controls for total HEX and HEXA activities. HEXB enzyme activity levels, in the form of the ratio of HEXA / total HEX, were not differential between sCAA and controls (p = 0.19) [**Figure 1 E**]. Correction for age (p = 0.20), total protein (p = 0.30), or age and total protein (p = 0.32) did not result in any significant differences between sCAA patients and controls.

CSF CTSB and CTSS levels were significantly associated (r_sp_ = 0.31, p = 0.02). Furthermore, significant associations were discovered between total HEX and CTSB (r_sp_ = 0.37, p = 0.005), and total HEX and CTSS (r_sp_ = 0.40, p = 0.003). Additionally, CTSS associated with HEXA (r_sp_ = 0.33, p = 0.01). Lastly, other associations which were discovered to be significant include CTSS and total protein (r_sp_ = 0.34, p = 0.01), total HEX and Aβ42 (r_sp_ = 0.26, p = 0.05), and total HEX and BCA (r_sp_ = 0.29, p = 0.03) [**Figure 2**].

**Figure 2:**
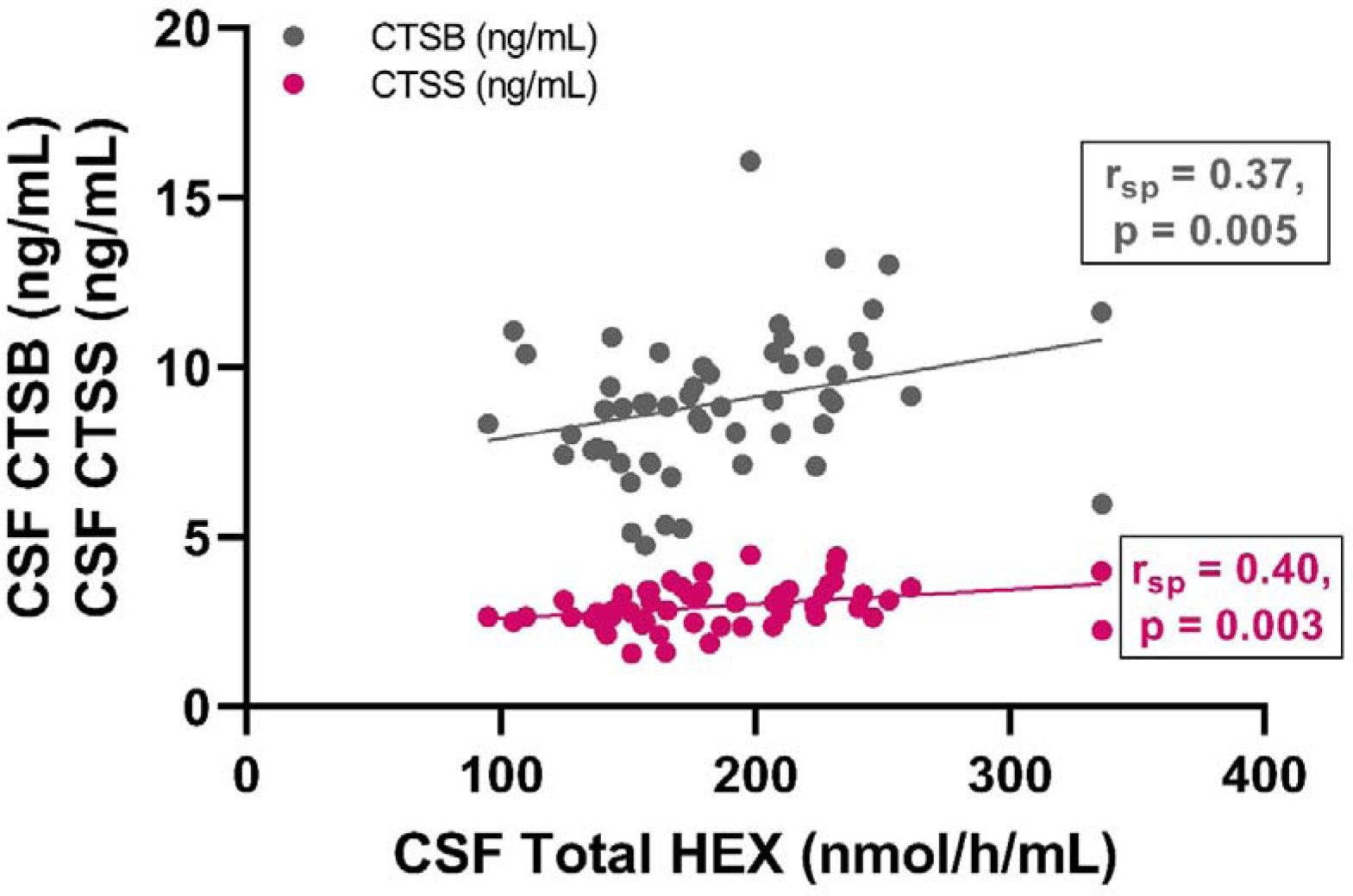
Scatter plot on the associations between CSF total HEX activity and CSF levels of CTSB/CTSS. Significant associations were discovered between total hexosaminidase activity and protein levels of CTSB (p = 0.005) and CTSS (p = 0.003). CTSB is shown in grey and CTSS is shown in pink. r_sp_ = Spearman correlation coefficient.

There were few significant associations between CTSB or CTSS, or total HEX or HEXB with radiological manifestations of sCAA. HEXA moderately, but significantly, associated with MOCA (r_sp_ = 0.27, p = 0.04), the number of lobar CMBs (r_sp_ = −0.30, p = 0.03), and CAA SVD score (r_sp_ = −0.34, p = 0.009). Stratification of the total population to respective diagnostic groups (CAA: r_sp_ = −0.25, p = 0.18/CON: r_sp_ = 0.05, p = 0.81) resulted in loss of significance of association between HEXA and CAA SVD score [**Figure 3**].

**Figure 3:**
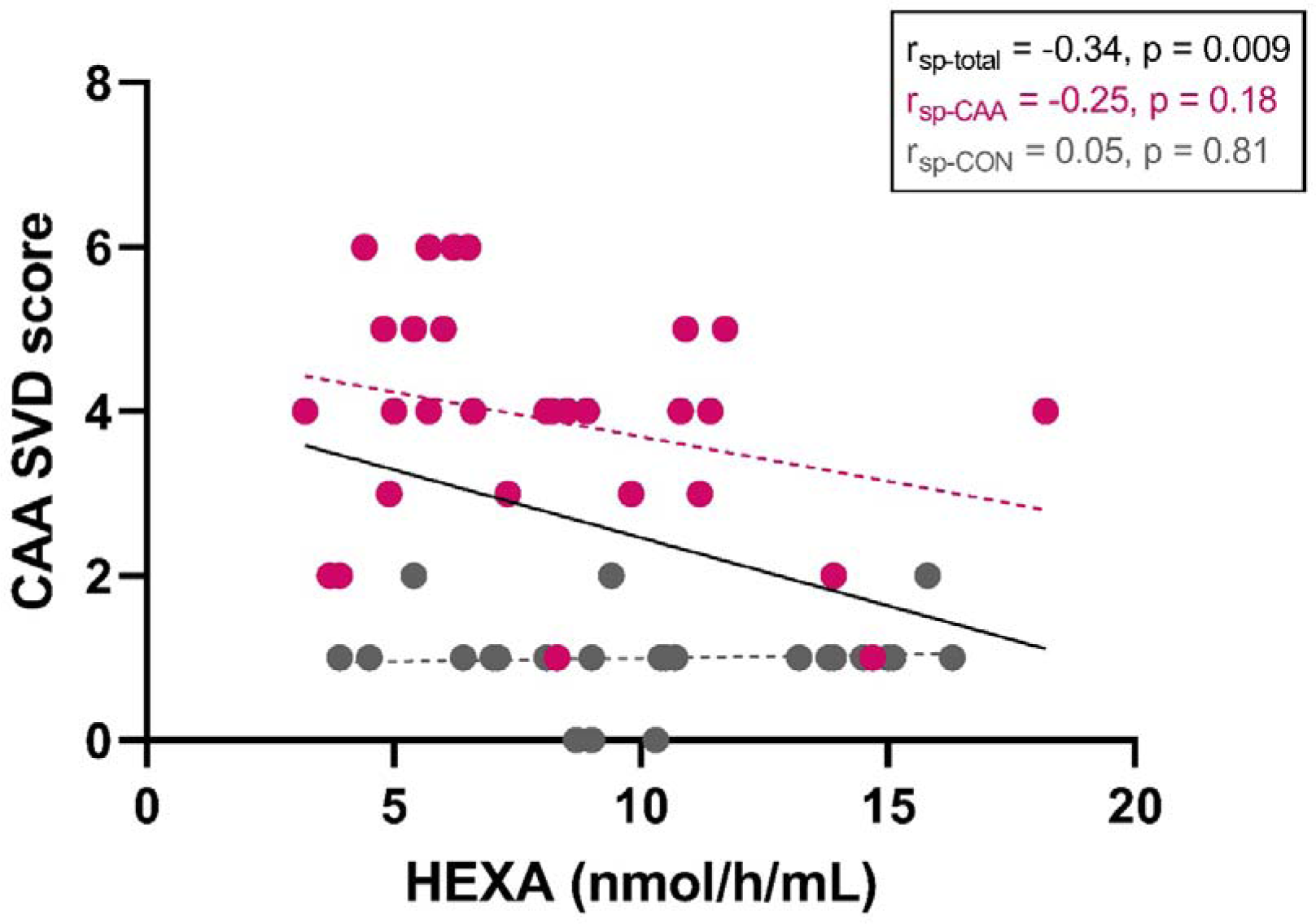
Correlations between CSF HEXA activity levels and CAA SVD score. Scatter plot on the association between CSF HEXA activity and composite CAA SVD score. Whereas for the total population (CAA + CON) a significant, negative association existed (p = 0.009), this significance disappeared after stratification for CAA diagnosis (CAA: p = 0.18; CON: p = 0.81). r_sp_ = Spearman correlation coefficient.

## Discussion

Our study has shown that the CSF CTSB and CTSS levels, and the CSF HEXB activity levels were not different between sCAA patients and controls. This is in contrast with earlier findings, which described increased expression of CTSB, CTSS and HEXB in CSF of rTg-DI rats compared to wild-type rats (5–8).

The observed discrepancy between findings in the CAA rat models and participants with CAA might be attributable to species-related disparities, as rats and humans exhibit significant biological distinctions, including variations in disease pathology, immune responses, and genetic factors (25). While animal models such as rTg-DI rats have been useful for research into pathogenesis and pathophysiology of CAA, they may not be able to fully replicate the complexity and heterogeneity of human sCAA (26). Another factor which might have affected the translatability of results between the rats and sCAA patients is the representativeness of the rTg-DI rat model for human sporadic CAA. The rTg-DI rat model presents with mostly capillary amyloid angiopathy, which in human CAA only occurs in CAA Type I (4). However, a majority of sCAA patients is known to suffer from a more macrovascular CAA Type II (arteries/arterioles), which raises questions on how well the rTg-DI rat is capable of modelling human sCAA pathology.

Another potential factor of influence on the discrepancy between results in rTg-DI rats and sCAA patients could be the distinction between protein levels and enzyme activity levels. Enzyme kinetics indicate that elevations in protein levels do not necessarily result in a proportionally increased enzyme activity and stable protein levels might nonetheless result in higher enzyme activity due to external circumstances. Our in-house CTSB/CTSS activity assays were capable of determining activity levels in human serum. While we based the design of our assays on successful assays described in literature, we were unable to sufficiently increase the analytical sensitivity for reliable analysis in CSF (22, 23). We have not been able to identify the reason behind the limited sensitivity of our in-house assays. Additionally, we were only able to approximate HEXB activity through an indirect method. Further comparison of CSF HEXA protease levels and activity rates might grant additional insight in the way in which differences in CSF protein expression and/or enzyme activity relate to each other in a translational sense between rats and humans, and whether this is affected by CAA pathology. Additionally, more sensitive (CTSB/CTSS) and more specific (HEXB) activity assays might allow for additional, essential improvements.

HEXA activity was decreased in sCAA patients compared to activity in controls, a difference which disappeared after correction for age and/or total protein. This is surprising, as earlier research had observed that HEXA was not differentially expressed between rTg-DI and WT rats. On the other hand, HEXB was differentially expressed in brain tissue/cerebrovascular tissue of rTg-DI rats. This might be due to the fact that the HEXA and HEXB isozymes have structural and functional similarities, but have distinct roles and substrate specificities (27).

Weak to moderate associations between HEXA activity and the MOCA score (positive) and number of microbleeds and CAA SVD score (negative) might support a relationship between HEXA activity and severity of CAA pathology. However, the robustness of these findings is limited, as significant associations were only found irrespective of sCAA diagnosis. Associations after stratification of sCAA diagnosis were not significant. The extent to which CAA pathology might induce inflammation or activate the immune system remains mostly unknown. However, severe perivascular neuroinflammation has been documented in both rTg-DI rats with advanced vascular amyloid pathology and human CAA patients moving from the non-haemorrhagic to the haemorrhagic phase of CAA (3, 28). It is possible that CAA pathology is capable of inducing increased expression and/or activities of lysosomal enzymes, including those of cathepsins and hexosaminidases. However, if and to which extent this occurs, is currently unknown.

CTSB and CTSS protein levels positively associated with each other and with total HEX activity levels, whereas CTSS also associated with HEXA activity levels. The basis for these associations are likely in the co-localization of respective proteins within lysosomes, which serve as a common environment for both cathepsins and hexosaminidases (29). This shared spatial proximity could potentially lead to coordination or correlation in their activities as part of lysosomal function. For example, cathepsins are known to play an active role in processes such as antigen presentation and immune responses, which can influence lysosomal function and through this affect the activity of lysosomal enzymes like HEXA and HEXB (30, 31).

The observed discrepancies between discovery results in rTg-DI rats and validation results in sCAA patients could be considered exemplary for the limited robustness of results in sCAA biomarker research (32). While challenging, these contrasting results offer valuable insights into the nuances of the disease and the intricacies of translating laboratory findings into clinical practice. Understanding these discrepancies is instrumental for enhancing the precision of biomarker translation, ensuring clinical relevance, and developing holistic biomarker panels that capture a broader spectrum of disease-related information. These insights, gained through the complexities of biomarker validation, can inform better-targeted treatment and intervention strategies, ultimately advancing our ability to diagnose, understand, and treat diseases like CAA.

## Conclusion

In conclusion, we were unable to validate previous observations, obtained in untargeted proteomics studies, on the enzymes CTSB, CTSS and HEXB as CSF biomarkers in sCAA. The observed discrepancies between a rat model of CAA and human patient samples underscore the complex interplay of factors influencing protein expression and enzyme activities. Future studies should aim to control for confounding (translational) variables and consider implications for better aligning preclinical models with human disease, ultimately advancing our ability to diagnose, treat, and understand CAA.

## Bibliography

1. Koemans EA, Chhatwal JP, van Veluw SJ, van Etten ES, van Osch MJP, van Walderveen MAA, et al. Progression of cerebral amyloid angiopathy: a pathophysiological framework. The Lancet Neurology. 2023;22(7):632–42.

2. Greenberg SM, Bacskai BJ, Hernandez-Guillamon M, Pruzin J, Sperling R, van Veluw SJ. Cerebral amyloid angiopathy and Alzheimer disease - one peptide, two pathways. Nat Rev Neurol. 2020;16(1):30–42.

3. Davis J, Xu F, Hatfield J, Lee H, Hoos MD, Popescu D, et al. A Novel Transgenic Rat Model of Robust Cerebral Microvascular Amyloid with Prominent Vasculopathy. The American Journal of Pathology. 2018;188(12):2877–89.

4. Thal DR, Ghebremedhin E, Rüb U, Yamaguchi H, Del Tredici K, Braak H. Two Types of Sporadic Cerebral Amyloid Angiopathy. Journal of Neuropathology & Experimental Neurology. 2002;61(3):282–93.

5. Vervuurt M, Schrader JM, de Kort AM, Kersten I, Wessels H, Klijn CJM, et al. Cerebrospinal fluid shotgun proteomics identifies distinct proteomic patterns in cerebral amyloid angiopathy rodent models and human patients. Acta Neuropathol Commun. 2024;12(1):6.

6. Schrader JM, Stanisavljevic A, Xu F, Van Nostrand WE. Distinct Brain Proteomic Signatures in Cerebral Small Vessel Disease Rat Models of Hypertension and Cerebral Amyloid Angiopathy. J Neuropathol Exp Neurol. 2022;81(9):731–45.

7. Schrader JM, Xu F, Van Nostrand WE. Distinct brain regional proteome changes in the rTg-DI rat model of cerebral amyloid angiopathy. Journal of Neurochemistry. 2021;159(2):273–91.

8. Schrader JM, Xu F, Lee H, Barlock B, Benveniste H, Van Nostrand WE. Emergent White Matter Degeneration in the rTg-DI Rat Model of Cerebral Amyloid Angiopathy Exhibits Unique Proteomic Changes. The American Journal of Pathology. 2022;192(3):426–40.

9. Yadati T, Houben T, Bitorina A, Shiri-Sverdlov R. The Ins and Outs of Cathepsins: Physiological Function and Role in Disease Management. Cells. 2020;9(7).

10. Xie Z, Meng J, Kong W, Wu Z, Lan F, Narengaowa, et al. Microglial cathepsin E plays a role in neuroinflammation and amyloid β production in Alzheimer’s disease. Aging Cell. 2022;21(3):e13565.

11. Pišlar A, Bolčina L, Kos J. New Insights into the Role of Cysteine Cathepsins in Neuroinflammation. Biomolecules. 2021;11(12).

12. Nakanishi H. Microglial cathepsin B as a key driver of inflammatory brain diseases and brain aging. Neural Regen Res. 2020;15(1):25–9.

13. Lowry JR, Klegeris A. Emerging roles of microglial cathepsins in neurodegenerative disease. Brain Res Bull. 2018;139:144–56.

14. Ni J, Lan F, Xu Y, Nakanishi H, Li X. Extralysosomal cathepsin B in central nervous system: Mechanisms and therapeutic implications. Brain Pathol. 2022;32(5):e13071.

15. Hou Y, Tse R, Mahuran DJ. Direct Determination of the Substrate Specificity of the α-Active Site in Heterodimeric β-Hexosaminidase A. Biochemistry. 1996;35(13):3963–9.

16. O’Brien JS. Five gangliosidoses. Lancet. 1969;2(7624):805.

17. Miyamoto E, Sato T, Matsubara T. Cyclization of Peptides Enhances the Inhibitory Activity against Ganglioside-Induced Aβ Fibril Formation. ACS Chem Neurosci. 2023;14(23):4199–207.

18. Linn J, Halpin A, Demaerel P, Ruhland J, Giese AD, Dichgans M, et al. Prevalence of superficial siderosis in patients with cerebral amyloid angiopathy. Neurology. 2010;74(17):1346–50.

19. Nasreddine ZS, Phillips NA, Bédirian V, Charbonneau S, Whitehead V, Collin I, et al. The Montreal Cognitive Assessment, MoCA: a brief screening tool for mild cognitive impairment. J Am Geriatr Soc. 2005;53(4):695-9.

20. De Kort AM, Kuiperij HB, Marques TM, Jäkel L, van den Berg E, Kersten I, et al. Decreased Cerebrospinal Fluid Amyloid β 38, 40, 42, and 43 Levels in Sporadic and Hereditary Cerebral Amyloid Angiopathy. Annals of Neurology. 2023;93(6):1173–86.

21. Charidimou A, Martinez-Ramirez S, Reijmer YD, Oliveira-Filho J, Lauer A, Roongpiboonsopit D, et al. Total Magnetic Resonance Imaging Burden of Small Vessel Disease in Cerebral Amyloid Angiopathy: An Imaging-Pathologic Study of Concept Validation. JAMA Neurol. 2016;73(8):994–1001.

22. Barrett AJ, Kirschke H. Cathepsin B, Cathepsin H, and cathepsin L. Methods Enzymol. 1981;80 Pt C:535–61.

23. Boutté AM, Hook V, Thangavelu B, Sarkis GA, Abbatiello BN, Hook G, et al. Penetrating Traumatic Brain Injury Triggers Dysregulation of Cathepsin B Protein Levels Independent of Cysteine Protease Activity in Brain and Cerebral Spinal Fluid. J Neurotrauma. 2020;37(13):1574–86.

24. Okada S, O’Brien JS. Tay-Sachs Disease: Generalized Absence of a Beta-D-N-Acetylhexosaminidase Component. Science. 1969;165(3894):698-700.

25. Zeiss CJ. From Reproducibility to Translation in Neurodegenerative Disease. ILAR Journal. 2017;58(1):106–14.

26. Jäkel L, Van Nostrand WE, Nicoll JAR, Werring DJ, Verbeek MM. Animal models of cerebral amyloid angiopathy. Clin Sci (Lond). 2017;131(19):2469–88.

27. Srivastava SK, Beutler E. Hexosaminidase-A and Hexosaminidase-B: Studies in Tay-Sachs’ and Sandhoff’s Disease. Nature. 1973;241(5390):463-.

28. Kozberg MG, Yi I, Freeze WM, Auger CA, Scherlek AA, Greenberg SM, et al. Blood–brain barrier leakage and perivascular inflammation in cerebral amyloid angiopathy. Brain Communications. 2022;4(5).

29. David A, Chazeirat T, Saidi A, Lalmanach G, Lecaille F. The Interplay of Glycosaminoglycans and Cysteine Cathepsins in Mucopolysaccharidosis. Biomedicines. 2023;11(3):810.

30. Riese RJ, Mitchell RN, Villadangos JA, Shi GP, Palmer JT, Karp ER, et al. Cathepsin S activity regulates antigen presentation and immunity. J Clin Invest. 1998;101(11):2351–63.

31. Man SM, Kanneganti TD. Regulation of lysosomal dynamics and autophagy by CTSB/cathepsin B. Autophagy. 2016;12(12):2504–5.

32. Erdogan BR, Michel MC. Building Robustness into Translational Research. In: Bespalov A, Michel MC, Steckler T, editors. Good Research Practice in Non-Clinical Pharmacology and Biomedicine. Cham: Springer International Publishing; 2020. p. 163–75.

